# A panel of biologically contained orthoebolaviruses for the screening of broad-spectrum antivirals

**DOI:** 10.64898/2026.07.28.741243

**Authors:** Jens Verlinden, Joren Stroobants, Jesus Sacramento Nonay, Charlotte Willems, Kato Govaerts, Bram Van Holm, Winston Chiu, Joost Schepers, Thibault Francken, Viktor Lemmens, Kurt Vermeire, Bert Vanmechelen

## Abstract

Filoviruses, particularly those of the *Orthoebolavirus* genus, pose ongoing public health threats due to their increasing frequency and geographic spread. However, research has been impeded by the biosafety level 4 classification of filoviruses. We previously reported the generation of biologically contained Ebola, Marburg and Sudan virus as lower biosafety level-compatible filovirus systems. In the present study, we expanded this repertoire to include biologically contained Taï Forest, Bundibugyo and Reston virus, thereby creating a near-complete toolkit for currently recognized human-relevant orthoebolaviruses. VP30-deficient viruses were generated with matching VP30 expressing cell lines. More specifically, we developed and optimized a dual-reporter system in VeroE6, Huh-7 and A549 cells, combining a virus-encoded enhanced green fluorescent protein reporter as readout for viral replication with a stably cellular-expressed nuclear mCherry marker for cytotoxicity assessment. Next, we screened two repurposing-oriented compound libraries comprising 640 small molecules against Taï Forest, Bundibugyo and Sudan virus, and subsequently cross-validated against Reston and Ebola virus. This approach identified multiple candidates with broad-spectrum activity across orthoebolaviruses, while also revealing virus-specific antivirals, with robust activity observed in both primate- and human-derived cell lines. Together, this work establishes a versatile and experimentally tractable lower biosafety level-compatible platform for the study of human-relevant orthoebolaviruses and the systematic discovery of broad-spectrum antiviral countermeasures against filoviruses.

## Introduction

Ebola virus (EBOV) and Sudan virus (SUDV), both members of the *Orthoebolavirus* genus and highly pathogenic to humans, were discovered in 1976 following two nearly simultaneous outbreaks in the Democratic Republic of the Congo and in South Sudan [1]. These early events were marked by high mortality rates and difficulties with containment, due to nosocomial transmission, reuse of contaminated medical equipment, and limited understanding of the causative agents [2, 3]. Since the 1990s, the frequency of confirmed orthoebolavirus outbreaks has risen sharply, with no five-year interval passing without at least one event [2]. In addition, four more orthoebolavirus species have been identified: Taï Forest virus (TAFV), Bundibugyo virus (BDBV), Reston virus (RESTV) and Bombali virus (BOMV), with the former two being pathogenic for humans. Taï Forest ebolavirus (TAFV) was detected in 1994 in Côte d’Ivoire following a single human infection linked to handling a chimpanzee carcass. Bundibugyo ebolavirus (BDBV) was first identified in Uganda in 2007, caused a second outbreak in the Democratic Republic of the Congo (DRC) in 2012, and is currently (2026) responsible for an ongoing Ebola disease outbreak in eastern DRC with cross-border spread to Uganda [4–7]. RESTV, discovered in non-human primates in the Philippines, is currently considered apathogenic to humans, however, asymptomatic RESTV-specific antibody responses have been detected in animal handlers and pig workers following exposure, suggesting a potential for human infection if viral evolution were to increase its human pathogenicity [8–10]. BOMV has only been observed in bats of the *Mops* genus [11, 12]. This rising trend in frequent outbreaks is also observed for the related Marburg virus (MARV), suggesting a broader increase in filovirus emergence [13]. Several ecological and socio-economic factors likely contribute to this rise, including climate change, deforestation, loss of bio-diversity and increased human–wildlife interaction, particularly via bushmeat consumption [14–17]. Moreover, the geographic range of filoviruses has proven broader than initially assumed. While EBOV and MARV outbreaks were historically confined to Central Africa, both have caused significant outbreaks in West Africa since 2014 [18, 19]. The 2014–2016 EBOV epidemic, originating in Guinea, evolved into the largest on record, affecting six neighboring African countries and highlighting the vulnerability of health systems unfamiliar with filoviruses [20]. Seroepidemiological and surveillance studies have since suggested potential co-circulation of multiple filoviruses in the same endemic regions, uncovering new hurdles for surveillance and outbreak response [21].

Small molecule antivirals (SMAs) emerge as an essential complementary approach to existing vaccination tools [22]. These compounds can be administered after symptom onset, are generally cost-effective, and often do not require cold-chain storage [23–25]. Furthermore, their utility is not always limited to a single virus species and can even be effective across virus families [26]. Considering the co-circulation of multiple filoviruses in endemic areas and the delay often associated with laboratory confirmation, partly due to frontline molecular tests being optimized for EBOV, timely etiological diagnosis is not always feasible. In this context, broad-spectrum antivirals could be deployed empirically in suspected cases, potentially improving early-stage management and outcomes. Nevertheless, very few small molecules have progressed through the antiviral pipeline for filoviruses. Remdesivir, a nucleotide analogue prodrug targeting the viral RNA polymerase, has shown *in vitro* and *in vivo* efficacy against EBOV and SUDV, but its performance in clinical trials (e.g., the PALM trial) was rather limited [23]. Similarly, favipiravir and brincidofovir showed early promise in preclinical studies. However, favipiravir failed to demonstrate clinical benefit in human trials, while brincidofovir yielded inconclusive results due to a limited sample size [27, 28]. The identification of such compounds is hampered by the biosafety level 4 (BSL-4) classification of filoviruses, which restricts research to a handful of laboratories worldwide [29]. Although reverse genetics techniques have been developed to circumvent these high-level requirements, each system has its limitations. For instance, minigenome systems and infectious virus-like particles (iVLPs) are limited to specific parts of the lifecycle or to single infectious rounds respectively, and rely on the co-transfection of viral protein-encoding plasmids [30]. In 2008, Halfmann *et al.* introduced a biologically contained EBOV system (EBOVΔVP30) by removing the essential transcription co-factor VP30 from the viral genome and supplying it *in trans* via engineered cell lines, enabling live virus replication in a restricted setting of VP30-expressing cells only [31]. Building on this concept, our group established similar systems for SUDV and MARV and demonstrated their value in robust antiviral screens for all three viruses, supporting the discovery of novel medical countermeasures against orthoebolaviruses [32–34].

In this study, we aim to expand the biologically contained filovirus platform with BDBVΔVP30, TAFVΔVP30 and RESTVΔVP30, having now a broadly representative toolkit of biologically contained alternatives for known human-infecting orthoebolaviruses. Using dual-reporter monoclonal cell lines and optimized screening workflows, we evaluated the antiviral activity of two small-molecule libraries provided by Medicines for Malaria Venture (MMV). By including EBOV hits from a prior biologically contained EBOV screen [34], and by using this previously established EBOV contained system and RESTVΔVP30 as confirmatory proxies, we show that this lower BSL-compatible multivirus platform is suitable for detecting broad-spectrum antivirals across orthoebolaviruses (Figure S1).

## Materials and methods

### Cell lines

African green monkey kidney cells (VeroE6; Vero C1008, American Type Culture Collection (ATCC)), human hepatocellular carcinoma cells (Huh-7; kindly provided by Ralf Bartenschlager, University of Heidelberg, Germany), adenocarcinomic human alveolar basal epithelial cells (A549, ATCC CCL-185) and human embryonic kidney epithelium 293FT cells (HEK293FT, ATCC) were grown in Dulbecco’s Modified Eagle Medium (DMEM; Thermo Fisher Scientific) supplemented with 10% Fetal Bovine Serum (FBS; Biowest, Nuaillé, France). HEK293FT cells were additionally maintained in the presence of 500 µg/ml Geneticin (G418; Thermo Fisher Scientific) according to the manufacturer’s recommendations. Except for HEK293FT cells, other cells were further supplemented with 2 mM L-glutamine (Thermo Fisher Scientific) and 8.8 mM sodium bicarbonate (Thermo Fisher Scientific). For Huh-7, the medium also contained 8.8 mM non-essential amino acids (NEAA, Thermo Fisher Scientific). As general antibiotics/antimycotic supplements, 1% Penicillin-streptomycin (Thermo Fisher Scientific), 0.2% Amphotericin B (Thermo Fisher Scientific) and 0.02% Gentamicin (Thermo Fisher Scientific) were used. Specific FBS concentrations and/or supplementary antibiotics used throughout experiments are indicated in the appropriate sections. All cell lines were maintained at 37 °C in a humidified incubator with 5% CO₂.

### Plasmids

Plasmids containing the full-length, VP30-deficient antigenome of either RESTV, BDBV or TAFV, in which the essential VP30 gene is replaced by an eGFP reporter cassette, were generated by a four-fragment assembly using the NEBuilder Hifi DNA assembly cloning kit (New England Biolabs (NEB)). The T7 backbone, including the T7 promoter and a C-terminal hepatitis delta virus ribozyme (HdVRz), used for the assembly of all three antigenomes was amplified from a T7 plasmid using the Q5 Hotstart-Fidelity 2X master mix (NEB) as described before [34]. For each virus, three ∼5600-7600-bp fragments were synthesized by Genscript Biotech in a pUC57 vector based on the reference genomes of the respective viruses (Genbank: NC_004161, NC_014373 and NC_014372). For RESTV and TAFV, these fragments contained flanking SmaI sites that were integrated during synthesis and used for excision from the pUC57 vector, whereas for the BDBV fragments we integrated SnaBI sites. The use and construction of pCAGGS plasmids encoding the EBOV nucleocapsid proteins has been described [34]. For RESTV, the coding sequences of the nucleocapsid genes were PCR-amplified from the assembled RESTV antigenome using Q5 Hot Start High-Fidelity 2× Master Mix (New England Biolabs). VP30 was derived from a separately synthesized fragment in a pUC57 vector (Genscript Biotech). The pCAGGS backbone was obtained via digestion of the previously described pCAGGS-L plasmid of EBOV [34]. Sequence verification of support plasmids was carried out with Sanger sequencing (Macrogen Europe, Amsterdam, The Netherlands). Sequence verification of antigenome constructs is described below.

### Lentiviral constructs

Cell lines stably expressing VP30 and/or mCherry were obtained by lentiviral transduction. The VP30 lentiviral constructs were made by insertion of the appropriate sequence into a pLenti6.3 vector (Thermo Fisher Scientific) with the NEBuilder HiFi DNA assembly cloning kit (NEB). The associated CMV promotor was substituted for a SFFV promotor to increase transcription efficiency. Additionally, an internal ribosomal entry site (IRES) cassette was placed between VP30 and the Blasticidin-S deaminase (bsd) gene by digestion of the pLenti6.3 vector with SpelI-HF and SalI-HF (NEB). Fragment ligation was realized with the Quick Ligation kit (NEB). A Kozak consensus sequence was placed upstream of the VP30 sequence to improve translation initiation. The mCherry lentiviral constructs were described previously [32].

### Lentiviral production and transduction

Lentivirus was produced by transfecting 50% confluent HEK293FT cells in T-25 flasks using Lipofectamine LTX & PLUS Reagent (Thermo Fisher Scientific). Briefly, transfection mixes included 3 μg of the VP30 lentiviral construct, 5.83 μg psPAX2, 3.17 μg pMD2.G, and 12 μl PLUS reagent in Opti-MEM (Thermo Fisher Scientific) and were mixed and incubated for 20 min before being added to the cells. Following 21 h incubation at 37 °C, 10 mM sodium butyrate was added for 3 h, after which medium was refreshed. Supernatant was harvested another 21 h later and centrifuged (2000 g, 15 min, 4 °C) to remove debris.

VeroE6, Huh-7, and A549 cells were transduced using the Virapower hiPerform T-Rex Gateway Expression System (Thermo Fisher Scientific) following the manufacturer’s instructions, with 6 μg/ml Polybrene (Sigma-Aldrich) added during transduction.

### Monoclonal selection

VeroE6, huh-7 and A549 cell lines stably expressing both mCherry and VP30 of either SUDV, RESTV, BDBV or TAFV were created by two separate lentiviral transductions. For SUDV, the VeroE6-VP30-mCherry monoclonal cell line generated in a previous manuscript was reused for relevant experiments here [32]. First, cells were transduced with a lentivirus containing the VP30 construct and put under the selection of Blasticidin (10μg/ml) followed by monoclonal selection. Next, a second transduction with a lentivirus containing the mCherry construct was carried out on the obtained monoclonal cell line and mCherry-positive cells were positively selected with the help of Geneticin (750 μg/ml). Monoclonal selection was carried out by limiting dilution as described previously [32]. Monoclonal cell lines derived from the same polyclonal cell line were compared in terms of cell growth and mCherry and eGFP-signal. Cells displaying aberrant cell morphology were removed from further experiments.

### Virus rescue

The rescue of SUDVΔVP30 and EBOVΔVP30 was described earlier [32, 34]. For the rescue of RESTV, BDBV and TAFV, 500,000 Huh-7 cells stably expressing EBOV VP30 (Huh-7-EBOV-VP30) were seeded in a 6-well plate and overnight incubated to reach 80-90% confluency [34]. Next, cells were transfected with a mixture containing 250 ng of the respective antigenome together with 125 ng EBOV NP, 125 ng EBOV VP35, 1000 ng EBOV L and 500 ng T7 polymerase, using jetOPTIMUS (Polyplus, Illkirch-Graffenstaden, France), according to the manufacturer’s protocol. For all viruses, 5 replicates and 1 negative control were included.

For BDBV and TAFV, rescued virus was transferred to freshly seeded Vero-BDBV-VP30 or Vero-TAFV-VP30 cells after respectively 4 and 6 days. Respectively 6 and 14 days after medium passaging, virus was purified by centrifuging the medium at 4500g for 15 minutes. Virus aliquots were stored at -80 °C until further use. For RESTV, cells were trypsinized at 14 days post transfection and 50% of the cell suspension was transferred to a new 6-well plate. This process was repeated three more times every seven days, after which medium was transferred to fresh Vero-RESTV-VP30 cells seven days after the last cell transfer. Virus was purified between days 19-33 after propagation on VeroE6 cells (depending on the isolate), when ∼100% eGFP-signal was reached.

### RNA extraction and Nanopore sequencing

RNA extraction was done using the QIAamp 96 Viral RNA kit (Qiagen, Venlo, The Netherlands), following the manufacturer’s protocol. Subsequently, the RNA was converted to cDNA and further amplified by Sequence-Independent Single Primer Amplification (SISPA) as described previously [35]. The amplified material was prepared for nanopore sequencing with the SQK-LSK109 kit (Oxford Nanopore Technologies (ONT), Oxford, UK) together with the EXP-NBD196 barcoding expansion (ONT). The library was then loaded on a R9.4.1 flow cell and sequenced on a GridION. Next, raw data was basecalled and barcodes were demultiplexed using the onboard software (ont-guppy-for-gridion v4.2.3). Read mapping against the according reference genome (Genbank: NC_004161, NC_014373 and NC_014372) was done using CLC workbench v22, whereafter Medaka v1.0.1 was used for consensus polishing and variant calling. Viral antigenome constructs were confirmed using the Rapid Barcoding Kit 24 V14 (ONT) following the manufacturer’s protocol. These libraries were also run on an R9.4.1 flow cell on the GridION and mapped against its reference plasmid sequences using CLC workbench as described.

### TCID50 determination

Double transduced monoclonal cells were seeded in 96-well plates in 100 μl medium and incubated (37 °C, 5% CO_2_). The next day, a ½ virus dilution series ranging from 1/12 to 1/3072 was made, and 50 μl of the dilution was transferred to infect the cells (12 replicates). High-content microscopic images were obtained for 10 consecutive days. Table S1 summarizes the specific conditions used in our assays depending on the virus and cell line. The EBOV assay on VeroE6 cells used as cross-validation was described before [34].

### Assay validation and high-content imaging

Validation of the assay was done by including remdesivir (GS-5734). Taking into consideration the conditions determined earlier, cells were seeded and incubated overnight in two 96-well plates. The next day, remdesivir was added to the cells in a ½ dilution series ranging from 50 or 10 (depending on the cell line) to 0.4, 0.02 or 0.0003 μM over 12 replicates (6 per plate) and incubated for one hour. Then, cells were infected with the according virus and incubated until read-out. Six negative (only medium) and positive (only virus) controls were included per plate. Outer wells were excluded due to observed edge-effects. Details about high-content imaging and robustness were described before [34]. The effect of CP-100356 on VeroE6 cells was evaluated in a similar way. CP-100356 is a high-affinity P-glycoprotein (P-gp) efflux pump inhibitor used to reduce drug export in VeroE6 cells [36]. 20,000 cells were seeded in a 96-well plate and incubated overnight. The next day, a dilution series of this efflux pump inhibitor was prepared and subjected to the cells in a range from 10 to 0.078 μM over 6 replicates. Read-out timing and additional information can be found in table S1.

### Antiviral screening assay

For the first round of screening, compounds spotted in 96-well plates as powder were kindly provided by Medicines for Malaria Venture (MMV), and subsequently dissolved in DMSO. SUDVΔVP30, BDBVΔVP30 and TAFVΔVP30 were chosen for this initial screening in VeroE6 cells, which were seeded one day prior to infection. The day of infection, intermediate four-fold dilutions of compounds were made in cell medium, in four concentrations ranging from 50 μM to 0.4 μM and added to the cells, followed by incubation. One hour later, virus was added to the cells. High-content images were obtained on an optimized read-out day specific for each virus. Assay-specific details and conditions can be found in table S1. Lay-out of the plates are illustrated in Figure S3.

### Hit confirmation and validation

Compound activity observed in our screening was confirmed using the initial DMSO solutions. For each compound to be confirmed against SUDVΔVP30, BDBVΔVP30 and TAFVΔVP30, an intermediate two-fold dilution of 8 concentrations was prepared over 2 replicates, allowing 3 compounds per plate. All other parameters were identical as described earlier.

For the compounds still displaying activity, fresh DMSO stocks were prepared from powder kindly provided by Evotech (Hamburg, Germany). Hit validation with these fresh stocks was carried out by testing each compound in two-fold dilutions over either 8 or 12 concentrations, depending on the compound, in triplicate. Compounds with the highest activity were selected to be tested as well in Huh-7 and A549 cells. Lay-out of the plates are illustrated in Figure S3.

### Data analysis and visualization

IC₅₀ and CC₅₀ values were determined by nonlinear regression using a four-parameter logistic (4PL) model in GraphPad Prism version 9.5.1 (GraphPad Software, San Diego, CA, USA), which was also used for generating dose–response curves. Venn diagrams were generated in a Jupyter Notebook environment using Python, with plots created using matplotlib (matplotlib.pyplot) and the venn package.

## Results

### Rescue and characterization of the biologically contained orthoebolaviruses BDBV, TAFV and RESTV

Based on the successful establishment of biologically contained EBOV, MARV and SUDV systems [33, 34], we sought to expand this toolkit to TAFV, BDBV and RESTV. For each virus, we generated a T7-driven VP30-deficient antigenome construct in which the essential VP30 gene was replaced by an eGFP reporter cassette, rendering viral replication dependent on VP30 supplied in trans. This virus-encoded eGFP provides an easy and direct readout of successful viral transcription and productive viral replication. In parallel, for each virus, VeroE6 and the human-derived Huh-7 and A549 cell lines were engineered to stably express the matching VP30 proteins. For rescue experiments, we used these VP30-only polyclonal lines, whereas VP30-mCherry monoclonal derivatives were generated only later for assay development. For the rescue of TAFVΔVP30, BDBVΔVP30 and RESTVΔVP30, VP30-expressing cells were transfected with the antigenome together with support plasmids encoding NP, VP35 and L from EBOV, as well as a T7 polymerase.

Using this approach, both TAFVΔVP30 and BDBVΔVP30 were rescued efficiently in Huh-7-EBOV-VP30 cells, as evidenced from a nearly complete eGFP-expressing, hence infected, cell monolayer within four and six days, respectively (data not shown). Subsequent propagation in the matching VeroE6-VP30 cells (by transfer of supernatant of the infected Huh-7 cells to freshly seeded VeroE6 cells) was also highly efficient, and resulted in a rapid increase in the number of infected VeroE6-VP30 cells within six and fourteen days for TAFVΔVP30 and BDBVΔVP30, respectively.

For RESTVΔVP30, initial rescue attempts in both VeroE6-RESTV-VP30 and Huh-7-RESTV-VP30 cells were unsuccessful, in line with previous reports describing the difficulties in generating both wild-type and chimeric RESTV using reverse genetics systems [37]. These studies have reported that the use of EBOV support plasmids, rather than homologous RESTV plasmids, can enhance rescue efficiency [37]. We explored this EBOV/RESTV combination strategy, but despite extensive efforts with different ratios of EBOV and RESTV support plasmids based on published protocols, none of these attempts were successful. Prompted by the findings of Deflubé et al., reporting that replication efficiency is influenced by terminal nucleotides [38], we re-evaluated the 3′ terminal sequence of the RESTV genome used for our antigenome design. Whereas the other orthoebolaviruses (EBOV, SUDV, BDBV and TAFV) contain a conserved GUCCA motif at the genome terminus, the RESTV reference sequence (NC_004161) ends in GUCC. After redesigning the RESTV antigenome construct with a terminal A nucleotide, we succeeded in rescuing RESTV, initially observed as slowly expanding clusters of eGFP-positive cells. However, in contrast to EBOVΔVP30, MARVΔVP30 and SUDVΔVP30 [33, 34], transfer of cell-free supernatant from these cultures to freshly seeded cells (either matching Huh-7-VP30 or VeroE6-VP30) did not result in detectable propagation. We therefore maintained the initially rescued Huh-7-EBOV-VP30 cell clusters by serial passaging of the infected cells until a near-complete eGFP positivity in the cell monolayer was obtained. Subsequent transfer of the supernatant yielded productive infection in VeroE6-RESTV-VP30 cells. Overall, RESTVΔVP30 rescue and propagation were markedly slower (19–33 days to reach maximal eGFP-positivity) as compared to the other biologically contained orthoebolaviruses.

Nanopore sequencing of the final virus stocks detected no non-synonymous mutations for BDBVΔVP30 and TAFVΔVP30, but two non-synonymous mutations were observed for RESTVΔVP30: D246N in VP40 and K755E in L.

### Assay optimization and validation

To establish antiviral screening assays for the biologically contained orthoebolaviruses, we generated monoclonal reporter cell lines for BDBV, TAFV and RESTV in VeroE6, Huh-7 and A549 cells, combining stable VP30 expression in the cytoplasm with a nuclear-localized mCherry reporter protein. Cell lines were generated by sequential lentiviral transduction and monoclonal selection, first for VP30 and subsequently for mCherry, to obtain clones with stable growth characteristics and robust mCherry expression. The optimized assay conditions and corresponding Z′-factors for each virus-cell line combination are summarized in table S1. With the exception of BDBV replication in Huh-7 cells (Z′ ≈ 0.5), all assays for BDBV, TAFV and RESTV showed high robustness, with most Z′-factors exceeding 0.9 [39], as evidenced by strong infection levels with limited variation across replicates (Figure 1). The same protocol was also applied to SUDV in Huh-7 and A549 cells, but these conditions did not provide sufficient and reproducible eGFP signals for downstream validation and were not pursued further.

**Figure 1:**
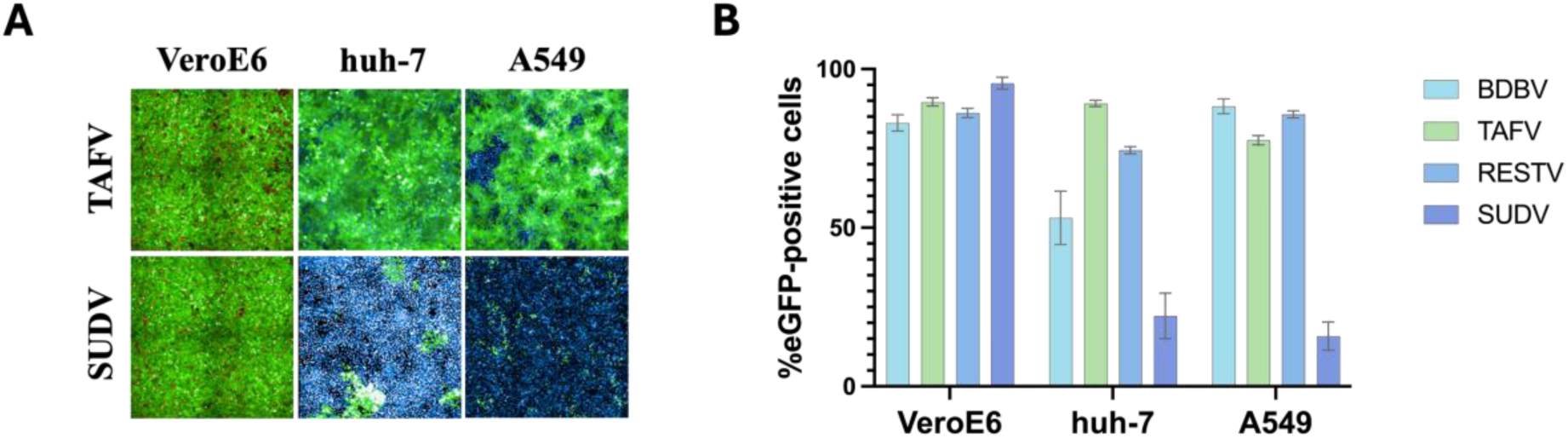
Replication efficiency of biologically contained orthoebolaviruses across different cell lines. High-content fluorescence images (panel A) and quantitative analysis (panel B) of VeroE6-VP30-mCherry, Huh-7-VP30, and A549-VP30 cells, each expressing the VP30 protein matching the corresponding biologically contained orthoebolavirus, infected under optimized conditions (Table S1). Images are shown for TAFV (top) and SUDV (bottom). Huh-7 and A549 cells were stained with 5 μM Hoechst 33342 nucleid acid stain to visualize and count nuclei. Replication efficiency is expressed as the percentage of eGFP-positive cells. Bars represent mean ± SD of 3-6 biological replicates.

In a next step we assessed the employability of our expanded ebolavirus toolkit for the screening of broad-spectrum antivirals. Remdesivir was used as a positive-control compound because of its established anti-filovirus activity [26, 40]. As expected, all assays displayed clear compound activity with high selectivity indices (Table 1 and Figure 2). When comparing IC_50_ values across the different viruses within the same cell line, values were generally consistent, typically differing by no more than three-fold. When comparing between cell lines for the same virus, remdesivir exhibited the greatest potency (lowest IC_50_ values and highest SI) in Huh-7 cells, followed by A549 cells, and then VeroE6 cells, highlighting the importance of having multiple orthogonal cell lines available for antiviral testing. IC_50_ values were noticeably higher for VeroE6 cells, in line with their known expression of P-glycoprotein transporters, which are capable of actively exporting xenobiotic compounds out of the cell [41]. Previously, we described the successful use of the P-glycoprotein inhibitor CP-100356 to address this limitation during the development and optimization of our SUDV assay, demonstrating enhanced antiviral efficiency of remdesivir [32]. However, for BDBV, the addition of CP-100356 resulted in reduced reporter gene expression, causing a marked decrease in assay robustness. To further characterize this interference, we conducted a dose-response analysis of CP-100356 against SUDVΔVP30, BDBVΔVP30, and TAFVΔVP30 using a two-fold dilution series ranging from 10 μM to 0.078 μM. This evaluation revealed a clear dose-dependent antiviral activity of CP-100356 against all three viruses, with some differences in potency across the three viruses. SUDV exhibited the lowest sensitivity (IC₅₀ = 4.4 μM), with no discernable effect at the 1 μM concentration previously used during assay development (Figure S2). Conversely, TAFVΔVP30 (2.8 µM) and BDBVΔVP30 (1.2 µM) were more sensitive and still displayed viral inhibition at the assay concentration (1 µM), explaining the reduced robustness of the VeroE6 assays in the presence of this compound. To prevent confounding effects on assay readout, CP-100356 was retained solely in the SUDV assay, given that antiviral activity was observed only at concentrations exceeding 3 µM, whereas no inhibitory effects were detected at the workable concentration.

**Figure 2:**
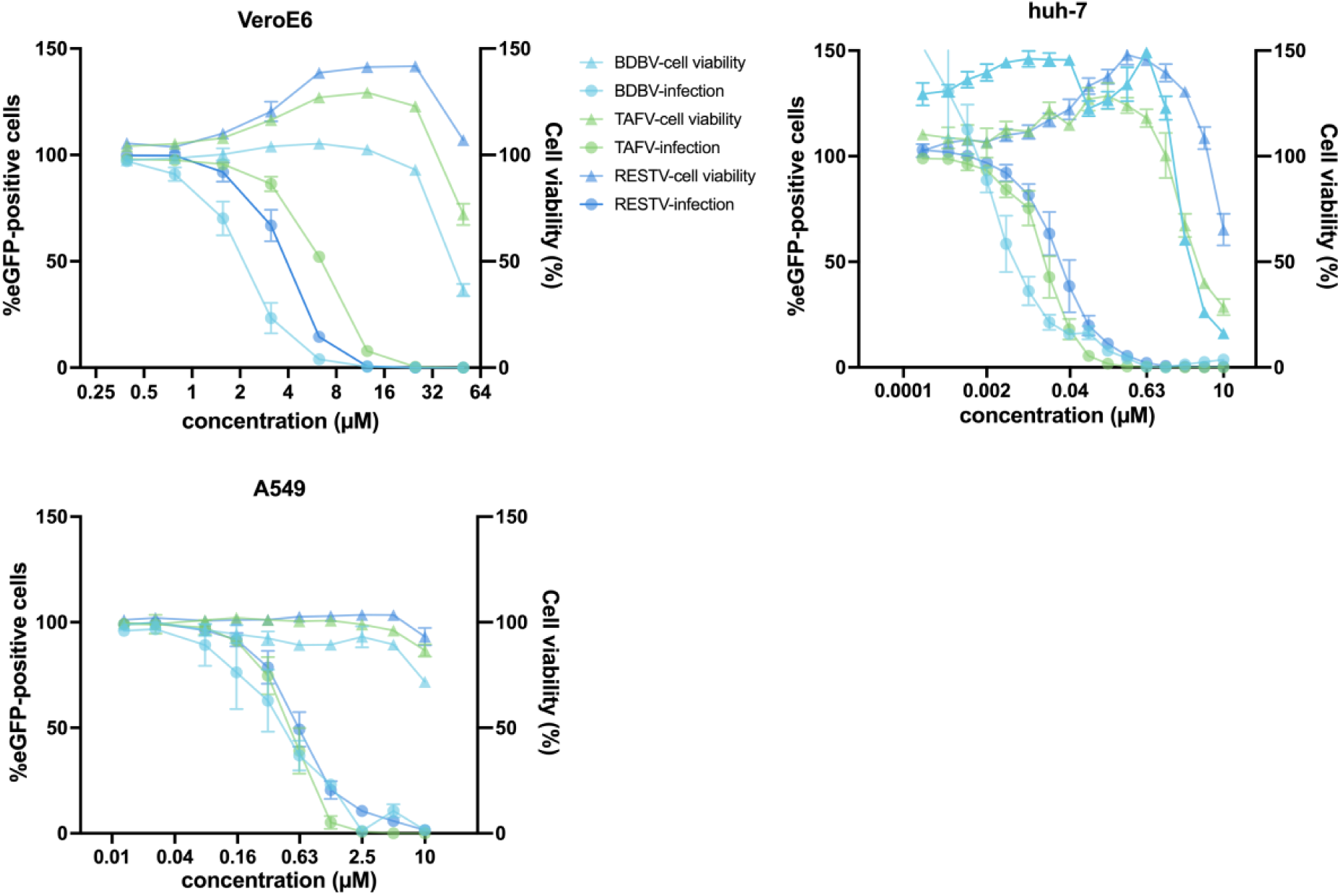
Validation of antiviral screening assays for BDBV, TAFV, and RESTV using remdesivir. Dose–response curves were generated for each virus–cell line combination using a 2-fold dilution series of remdesivir ranging from 50 μM to 0.39 μM (VeroE6) or lower where needed (Huh-7, A549). Cell lines stably expressed the VP30 protein corresponding to the biologically contained virus tested, together with nuclear mCherry. Antiviral activity was quantified by the percentage of eGFP-positive cells (circles, left y-axis) while cell viability was assessed through normalized mCherry read-out (triangles, right y-axis). Decreases in cell viability were interpreted as compound-induced cytotoxicity. Cell viability values above 100% result from normalization to virus-infected controls exhibiting low-level cytopathic effects, which are reduced upon inhibition of viral replication by remdesivir at non-cytotoxic concentrations. Each condition was tested in 3-6 replicates across two independent plates. Data are presented as mean ± standard deviation. Figures were generated using GraphPad Prism.

**Table 1:** Remdesivir performance across virus–cell line combinations.

| **Virus** | **Cell line\*** | **IC<sub>50</sub>** | **CC<sub>50</sub>** | **SI** |
| --- | --- | --- | --- | --- |
| BDBV | VeroE6 | 2.33 | 44.82 | 19.28 |
|  | Huh-7 | 0.006 | 3.36 | 560 |
|  | A549 | 0.43 | > 10 | > 23.41 |
| TAFV | VeroE6 | 6.55 | > 50 | > 7.63 |
|  | Huh-7 | 0.015 | 4.48 | 298 |
|  | A549 | 0.49 | > 10 | > 20.24 |
| RESTV | VeroE6 | 4.13 | > 50 | > 12.10 |
|  | Huh-7 | 0.028 | > 10 | > 357 |
|  | A549 | 0.62 | > 10 | > 16.07 |
| SUDV | VeroE6 | 2.33 | > 50 | > 21.47 |

\*each expressing a VP30 protein matching the respective virus.

### Screening of compound libraries for SUDV, BDBV and TAFV

Next, we screened the same MMV Pandemic Response Box together with the more recent Global Health Priority Box (240 compounds), totaling 640 small molecules with known or predicted antimicrobial activity. This approach enabled direct comparison between historical EBOVΔVP30 data and newly generated results [34]. Due to limited compound availability, the three human-pathogenic orthoebolaviruses (i.e., BDBV, TAFV and SUDV) were included in the primary screening phase, while RESTVΔVP30 was subsequently employed in downstream cross-validation steps [32, 42]. From the prior EBOVΔVP30 screen [34], the four most active CD3 (Centre for Drug Design and Discovery) compounds (Z-FA-FMK, dalbavancin, benproperine, and Evans Blue) were also included for cross-validation. Figure S3 illustrates the general screening and validation workflow.

### Compound screening and hit confirmation

For the screening, each compound was evaluated at four concentrations in a four-fold dilution series starting at 50 μM (50 – 12.5 – 3.13 – 0.78 µM) on VeroE6 cells. For each compound and dilution, the percentage of eGFP-positive cells and the total number of cells were normalized to virus-infected, untreated control conditions. Compounds were selected for follow-up testing when the reduction in infection (%eGFP-positive cells) exceeded the reduction in cell viability by more than 40 percentage points either at both of the two lowest tested concentrations, or at least at three of the four concentrations tested. Based on the criteria, 16, 26, and 21 compounds were selected for follow-up testing against SUDVΔVP30, BDBVΔVP30 and TAFVΔVP30, respectively. As indicated in Figure 3, multiple of these compounds had overlapping antiviral activity across two or all three viruses. For the confirmation step, compounds were tested in duplicate in a two-fold dilution series ranging from 50 μM to 0.39 μM to rule out false positive results.

**Figure 3:**
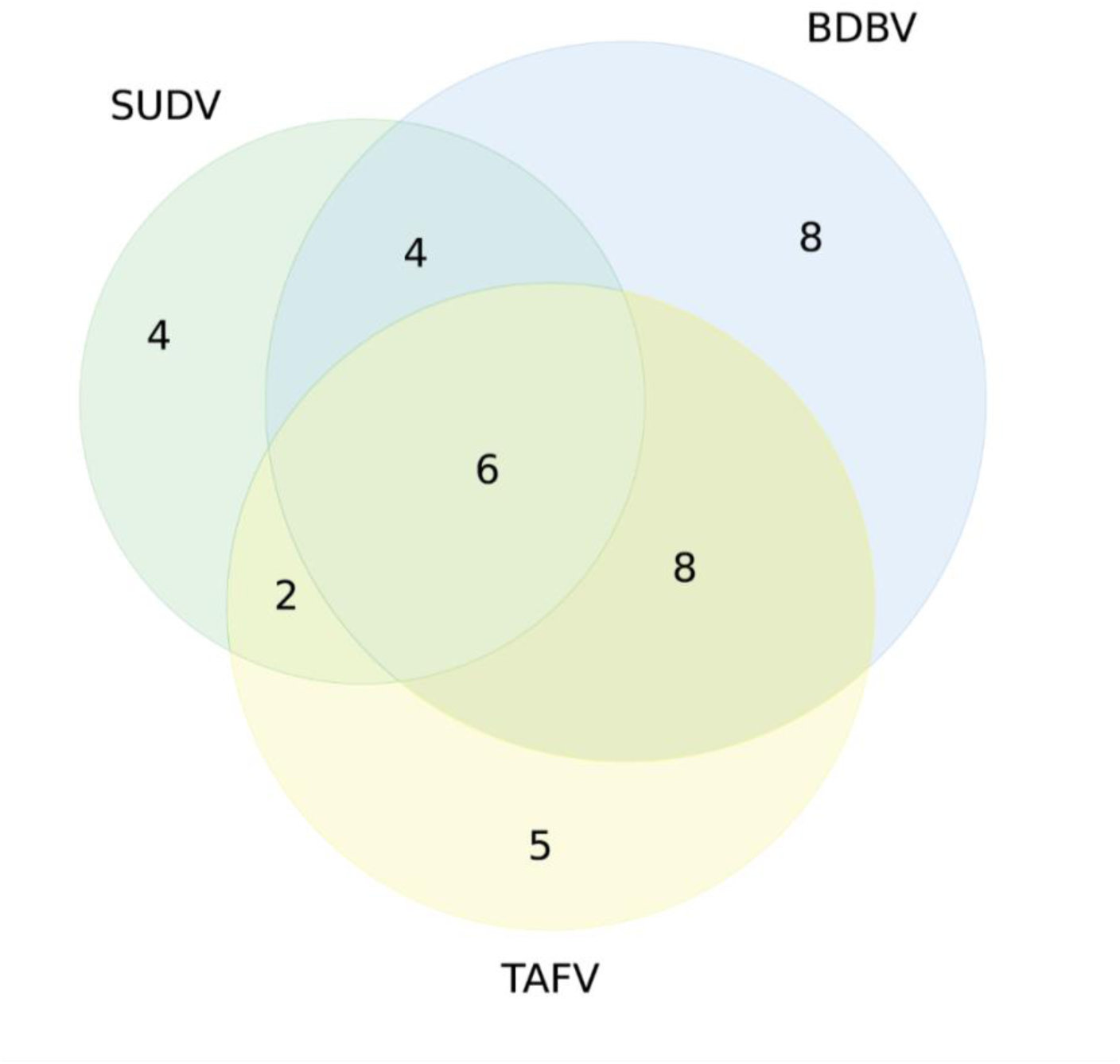
Overlap of antiviral hits identified in the primary screening against SUDV, BDBV, and TAFV in VeroE6 cells. Venn diagram illustrating the distribution of compounds showing antiviral activity in the initial screening. Numbers represent compounds meeting the predefined activity threshold, defined as at least a 40% difference between antiviral inhibition and the reduction in cell viability. Overlapping regions indicate compounds active against two or all three viruses.

### Hit validation

For compounds displaying selectivity indices (SI) ≥ 7, fresh powder stocks were procured to properly assess antiviral activity and to cross-validate against all other viruses in our toolkit, including EBOVΔVP30 [34]. An overview of these 15 compound hits in VeroE6 cells, including IC₅₀, CC₅₀, and SI-values, is presented in Table 2. Among the tested compounds, Z-FA-FMK and LHVS exhibited broad-spectrum activity against all five viruses, as illustrated in Figure 4. Several additional candidates showed activity against at least four viruses, including CHEMBL93139, MMV1795510, MMV893278, and MMV674884, while others displayed more selective antiviral profiles. Dalbavancin, Benproperin and Evans blue, coming from a prior study, failed to reach selectivity thresholds, as opposed to Z-FA-FMK, and are therefore not further included.

**Figure 4:**
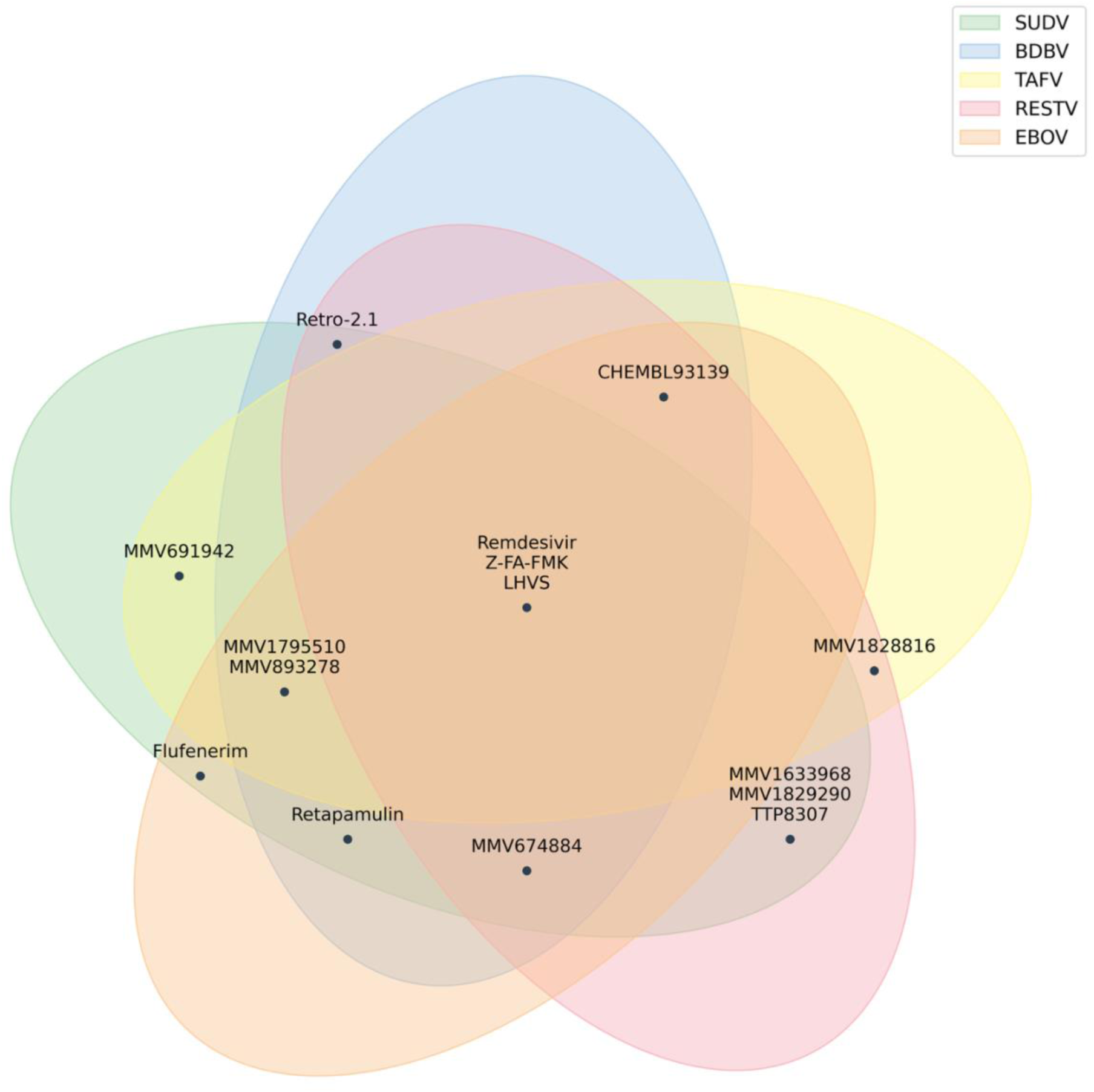
Overview of antiviral hit compounds identified in VeroE6 cells. Venn diagram illustrating compounds exhibiting antiviral activity against biologically contained SUDV, BDBV, TAFV, RESTV and EBOV. Compounds were considered active when displaying a SI greater than 7. Overlapping regions represent compounds active against multiple viruses, highlighting broad-spectrum candidates.

**Table 2:** Antiviral activity, cytotoxicity and selectivity indices of top hit compounds against SUDVΔVP30, BDBVΔVP30, TAFVΔVP30, RESTVΔVP30, and EBOVΔVP30 on VeroE6 cells.

| Compound | Trivial name | SUDVΔVP30 |  |  | BDBVΔVP30 |  |  | TAFVΔVP30 |  |  | RESTVΔVP30 |  |  | EBOVΔVP30 |  |  |
| --- | --- | --- | --- | --- | --- | --- | --- | --- | --- | --- | --- | --- | --- | --- | --- | --- |
|  |  | IC<sub>50</sub><br>(μM) | CC<sub>50</sub><br>(μM) | SI<sup>°</sup> | IC<sub>50</sub><br>(μM) | CC<sub>50</sub><br>(μM) | SI<sup>°</sup> | IC<sub>50</sub><br>(μM) | CC<sub>50</sub><br>(μM) | SI<sup>°</sup> | IC<sub>50</sub><br>(μM) | CC<sub>50</sub><br>(μM) | SI<sup>°</sup> | IC<sub>50</sub><br>(μM) | CC<sub>50</sub><br>(μM) | SI<sup>°</sup> |
| — | Z-FA-FMK | 1.6 | >50 | >**32.2** | <0.4 | >50 | >**128** | 2.3 | >50 | >**21.3** | 3.1 | >50 | >**15.9** | 2.3 | >50 | >**21.8\*** |
| MMV1804275 | LHVS | <0.8 | >50 | **64** | <0.4 | 40.8 | **104.5** | <0.4 | 24.4 | >**62.5** | 1.3 | 25.9 | **20.4** | 1.9 | >50 | >**26.5** |
| MMV893278 | — | 5.8 | >50 | >**8.6** | 1.2 | >50 | >**43.5** | 1.3 | >50 | >**38.3** | 38 | >50 | 1.3 | 3 | >50 | >**17** |
| MMV1633674 | Retapamulin | 2.3 | 32.6 | **14.1** | 1.3 | 32.1 | **24.1** | 8.5 | 27.9 | 3.3 | 7.9 | 47.2 | 6 | 3.7 | >50 | >**13.4\*** |
| MMV1782214 | CHEMBL93139 | 2.1 | 8.2 | 3.9 | 1.8 | 33.4 | **18.6** | 1.8 | 18.4 | **10.4** | 2.5 | 31.8 | **12.5** | 1.3 | 36.4 | **27.9** |
| MMV674884 | — | 4.1 | >50 | >**12.2** | 1.8 | >50 | >**28.2** | 2.1 | 6.1 | 2.9 | 5.2 | >50 | >**9.6** | 2.9 | >50 | >**17.2** |
| MMV1582492 | Retro-2.1 | <0.8 | >100 | >**128** | <0.7 | >50 | >**72.2** | <0.4 | 0.6 | >1.5 | 3.7 | >50 | >**13.4** | >50 | >50 | / |
| MMV1795510 **\$** | — | 1.3 | 9.8 | **7.5** | <0.4 | 8.7 | **22.3** | 0.6 | 5 | **7.9** | 4.3 | 12.5 | 2.9 | 0.7 | 7.2 | **10.9** |
| MMV1633968 | — | 3 | 92.6 | >**30.6** | >50 | >50 | / | >50 | >50 | / | 3 | >50 | >**16.5** | >50 | >50 | / |
| MMV691942 | — | 2.6 | >50 | >**19** | >50 | >50 | / | 0.6 | 10.2 | **17.7** | >50 | >50 | / | >50 | >50 | / |
| MMV1794206 | Flufenarim | <0.8 | 36.9 | **47.3** | >50 | >50 | / | >50 | >50 | / | 8.2 | 32.6 | 4 | 0.5 | 44.6 | **100.1** |
| MMV1782211 | TTP8307 | 5.5 | >50 | >**9.1** | 34.2 | >50 | >1.5 | 2.8 | 6.8 | 2.4 | 2.6 | >50 | >**19.1** | >50 | >50 | / |
| MMV1829290 | — | 6.3 | >50 | >**8** | >50 | >50 | / | >50 | >50 | / | 6.3 | >50 | >**8** | 26.9 | >50 | >1.9 |
| MMV1828816 | — | >50 | >50 | / | 9 | 22.8 | 2.5 | 8.4 | 17.6 | 2.1 | 5.1 | 21.3 | 4.2 | >50 | >50 | / |
| MMV1577456 | Bifentrin | >50 | >50 | / | 7.8 | >50 | >6.5 | 8.4 | 41.1 | 4.9 | 11.4 | >50 | >4.4 | >50 | >50 | / |

<sup>°</sup>SI>7 marked in bold.
\* Presented in a previous publication [34].
**\$** Insufficient compound material available for orthogonal validation.

While the overall antiviral profiles were largely consistent across assays, a few unexpected discrepancies were observed. For instance, Retro-2.1 displayed marked cytotoxicity in the TAFV assay, despite showing no apparent toxicity in the corresponding assays of BDBV and RESTV. This discrepancy may reflect differences between individual monoclonal cell lines although this was not investigated further. Compounds having activity against at least three viruses in VeroE6 cells were subsequently validated in two human-derived cell lines, huh-7 and A549 against BDBVΔVP30, TAFVΔVP30 and RESTVΔVP30 to assess antiviral efficacy and cytotoxicity across distinct cellular backgrounds and thereby strengthen the translational relevance of the screening results.

Upon orthogonal validation of selected compounds in human-derived cell lines against BDBVΔVP30, TAFVΔVP30, and RESTVΔVP30, the previously identified broad-spectrum antivirals Z-FA-FMK and LHVS retained strong activity across all viruses tested (Table 3). In addition, MMV893278 and retapamulin exhibited potent inhibitory effects against both BDBVΔVP30 and RESTVΔVP30. In contrast, compounds such as MMV674884, CHEMBL93139, and Retro-2.1, which had shown promising activity in VeroE6 cells, displayed markedly reduced selectivity in human cell lines, with antiviral activity observed only in isolated instances. This lack of reproducibility across cell lines underscores the importance of orthogonal validation in physiologically relevant models to confirm compound efficacy and to prevent overestimation of antiviral potential based solely on VeroE6 cell-based assays. Akin to testing in VeroE6 cells, compound toxicity profiles showed in some cases strong variation across different virus-cell line combinations.

**Table 3:**
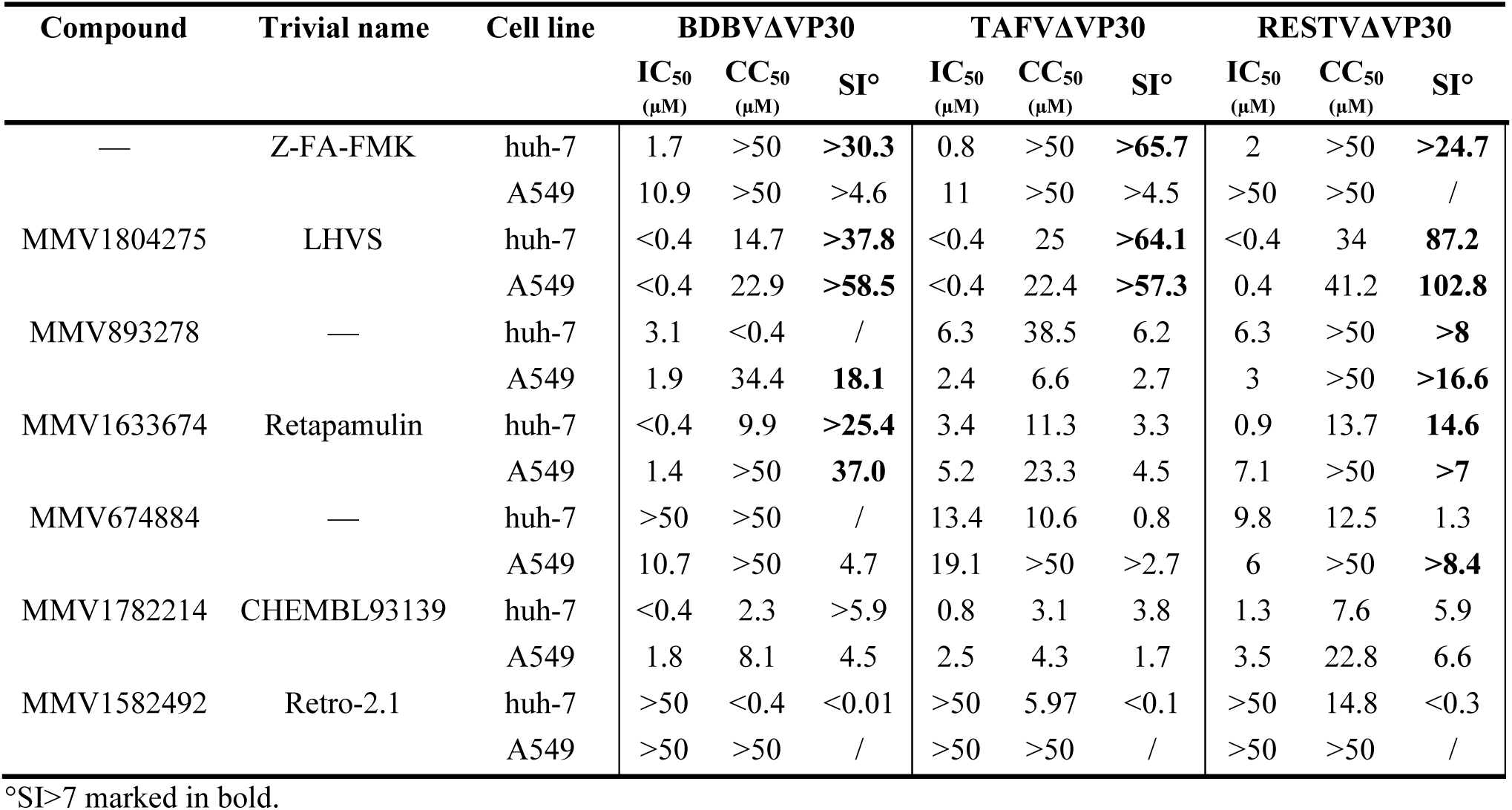
Orthogonal validation of selected hit compounds in human cell lines Huh-7 and A549.

| Compound | Trivial name | Cell line | BDBVΔVP30 |  |  | TAFVΔVP30 |  |  | RESTVΔVP30 |  |  |
| --- | --- | --- | --- | --- | --- | --- | --- | --- | --- | --- | --- |
|  |  |  | IC<sub>50</sub><br>(μM) | CC<sub>50</sub><br>(μM) | SI° | IC<sub>50</sub><br>(μM) | CC<sub>50</sub><br>(μM) | SI° | IC<sub>50</sub><br>(μM) | CC<sub>50</sub><br>(μM) | SI° |
| — | Z-FA-FMK | huh-7 | 1.7 | >50 | >**30.3** | 0.8 | >50 | >**65.7** | 2 | >50 | >**24.7** |
|  |  | A549 | 10.9 | >50 | >4.6 | 11 | >50 | >4.5 | >50 | >50 | / |
| MMV1804275 | LHVS | huh-7 | <0.4 | 14.7 | >**37.8** | <0.4 | 25 | >**64.1** | <0.4 | 34 | **87.2** |
|  |  | A549 | <0.4 | 22.9 | >**58.5** | <0.4 | 22.4 | >**57.3** | 0.4 | 41.2 | **102.8** |
| MMV893278 | — | huh-7 | 3.1 | <0.4 | / | 6.3 | 38.5 | 6.2 | 6.3 | >50 | >**8** |
|  |  | A549 | 1.9 | 34.4 | **18.1** | 2.4 | 6.6 | 2.7 | 3 | >50 | >**16.6** |
| MMV1633674 | Retapamulin | huh-7 | <0.4 | 9.9 | >**25.4** | 3.4 | 11.3 | 3.3 | 0.9 | 13.7 | **14.6** |
|  |  | A549 | 1.4 | >50 | **37.0** | 5.2 | 23.3 | 4.5 | 7.1 | >50 | >**7** |
| MMV674884 | — | huh-7 | >50 | >50 | / | 13.4 | 10.6 | 0.8 | 9.8 | 12.5 | 1.3 |
|  |  | A549 | 10.7 | >50 | 4.7 | 19.1 | >50 | >2.7 | 6 | >50 | >**8.4** |
| MMV1782214 | ChEMBL93139 | huh-7 | <0.4 | 2.3 | >5.9 | 0.8 | 3.1 | 3.8 | 1.3 | 7.6 | 5.9 |
|  |  | A549 | 1.8 | 8.1 | 4.5 | 2.5 | 4.3 | 1.7 | 3.5 | 22.8 | 6.6 |
| MMV1582492 | Retro-2.1 | huh-7 | >50 | <0.4 | <0.01 | >50 | 5.97 | <0.1 | >50 | 14.8 | <0.3 |
|  |  | A549 | >50 | >50 | / | >50 | >50 | / | >50 | >50 | / |

°SI>7 marked in bold.

### Global activity profile of validated hit compounds

To provide an intuitive and comparative overview of compound performance, validated hit compounds were summarized across all tested viruses and cell lines using a categorical activity classification (Figure 5). For each virus–cell line combination, compounds were considered effective when exhibiting a selectivity index greater than 7. This integrative representation in performance-based ranked order enables rapid visual prioritization of compounds based on both breadth of antiviral activity and cross-cell line robustness, rather than on individual IC₅₀ values alone.

**Figure 5:**
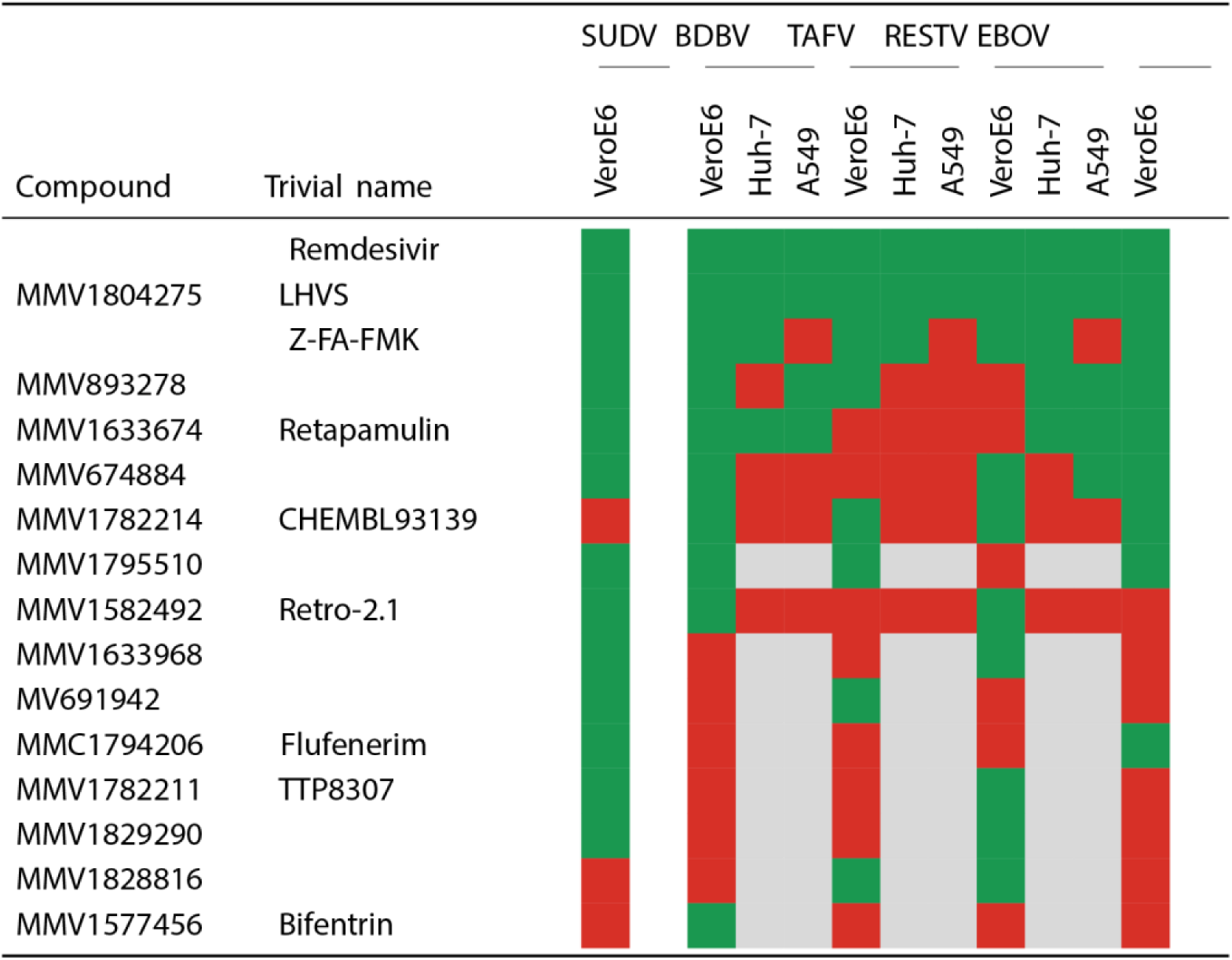
Antiviral hit performance across orthoebolaviruses and cell lines. Validated compounds are summarized according to their antiviral activity against biologically contained SUDV, BDBV, TAFV, RESTV, and EBOV across Vero E6, Huh-7, and A549 cell lines. Compound efficacy was classified based on a SI threshold of >7, with effective combinations indicated in green and inactive combinations in red. Grey cells indicate compounds that were not validated in human cell lines.

## Discussion

Small-molecule antivirals are particularly attractive in outbreak settings because they can be administered after symptom onset and are generally easier to deploy than vaccines or antibodies [22, 25, 43–45]. These compounds can be broadly divided into direct-acting antivirals (DAAs), which target viral proteins, and host-directed antivirals (HDAs), which target host factors or pathways required for the viral life cycle. While DAAs are susceptible to resistance mutations, HDAs tend to be more robust in this regard, though they may carry a higher risk of off-target effects [46, 47]. Combination therapies integrating both groups with antibodies and vaccines may therefore offer the most balanced approach to outbreak preparedness [47, 48]. Notably, SMAs displaying broad-spectrum activity are highly valuable in contexts where multiple viruses co-circulate and diagnostic capabilities are delayed [48]. Despite their promise, only a handful of broad-spectrum compounds are in the therapeutic pipeline, though with varying clinical relevance [23, 27, 49, 50].

A major bottleneck in filovirus research remains their classification as risk group 4 pathogens, which requires handling in biosafety level-4 (BSL-4) laboratories and continues to restrict antiviral discovery and therapeutic development [30, 33]. Biologically contained filoviruses provide a practical lower biosafety alternative [31], and our group previously established such systems for EBOV, SUDV, and MARV [33, 34]. In the present study, we extended this platform to BDBV, TAFV, and RESTV, thereby having a broadly representative toolkit of known human-relevant orthoebolaviruses. Whereas BDBVΔVP30 and TAFVΔVP30 could be incorporated readily, RESTVΔVP30 proved substantially more difficult to rescue and propagate, resembling the stringent requirements observed for MARVΔVP30 [33]. Efficient propagation on VeroE6 cells, as the gold standard cell line for filovirus propagation [51, 52], was not achieved by direct supernatant transfer and required alternative approaches. We suspect that either subthreshold viral doses or suboptimal cellular VP30 expression hindered initial propagation.

A key innovation of our platform was a dual-reporter design combining virus-encoded eGFP and nuclear-localized mCherry read-outs as proxies for viral replication and toxicity, respectively, extending described SARS-CoV-2 screening strategies [53]. Assay conditions were optimized to induce minimal CPE at the selected endpoint. This allowed both parameters to be quantified in the same infected wells by normalization to virus-infected controls, preserving a biologically relevant measure of compound effects in infected cells. An additional advantage of this format is its suitability for high-throughput screening, as it reduces material requirements compared with parallel antiviral and cytotoxicity plates. Follow-up testing with traditional approaches can, however, be readily performed for prioritized hits when needed.

VeroE6 cells were used as the primary screening platform because their flat, monolayer growth is well suited for high-content imaging [54], whereas Huh-7 and A549 cells are more demanding to culture and more difficult to analyze quantitatively due to multilayer growth [55]. Nevertheless, the robust performance of the human-derived assays supported their use for orthogonal hit validation, which is important because VeroE6 cells do not fully reflect human cellular biology [51]. Cell-type-specific host factor expression can influence both viral replication and the activity of HDAs [47], underscoring the value of validating hits in human-derived cell lines to improve translational relevance [56]. Furthermore, the use of monoclonal reporter cell lines improves long-term stability and reproducibility but also introduces potential limitations, as random lentiviral integration can generate clone-specific genetic alterations, including in genes that affect compound metabolism or viral replication [57]. As a result, antiviral and toxic effects of certain compounds may be under- or overestimated. This may have contributed to isolated discrepancies in compound behavior, such as the marked cytotoxicity of Retro-2.1 in the TAFV VeroE6 assay that was not reproduced in corresponding assays.

Examination of our hit set coming from our screening design indicates that not all compounds have the same scientific or translational meaning. Broadly, the hits can be categorized into four groups. A first group consists of tool compounds that serve primarily as benchmarks and proof-of-principle. Z-FA-FMK, previously active against EBOV [34], and LHVS exemplify this category: both showed the broadest antiviral activity, including in human-derived cell lines, but are cathepsin inhibitors targeting host enzymes with multiple cellular functions [58, 59], making systemic use prone to toxicity and tolerability issues. Their identification nonetheless underscores the validity of our assays and highlights their continued value as experimental tools for probing filovirus entry. A second category comprises hits that are likely limited by toxicity or pharmacology. MMV1795510 and CHEMBL93139, for example, displayed apparent broad activity, but also showed consistent cytotoxicity, making them unattractive development candidates despite their breadth. Finally, a third category encompasses genuine development candidates. Retapamulin and MMV893278 stand out in this category. Retapamulin is an approved topical antibiotic [60], but its pleuromutilin scaffold could form the basis for novel antiviral derivatives. MMV893278 currently lacks prior documentation in the antiviral field, yet retained activity with favourable selectivity in human-derived cell lines, making it an attractive starting point for further mechanistic work. Both compounds offer promising scaffolds for medicinal chemistry optimization to enhance antiviral potency and pharmacokinetic properties. Altogether, our findings also highlight the strength of repurposing-oriented libraries, which often contain molecules with characterized pharmacology and toxicology, thereby enabling more rapid downstream development compared to de novo screening [47, 61].

In addition to broad-spectrum candidates, we identified compounds with more restricted antiviral profiles. This reinforces that activity does not necessarily extend uniformly across orthoebolaviruses and highlights the importance of evaluating multiple species rather than relying on EBOV alone as prototype species [62]. At the same time, some observed differences may reflect clone-specific effects associated with the use of monoclonal cell lines [57].

At the time of writing, the current BDBV outbreak in the DRC and Uganda underscores the vulnerabilities highlighted throughout this manuscript [4]. Although the precise spillover event remains unresolved, retrospective modelling suggests that the index transmission likely occurred in mid-to-late February [63], after which severe disease accumulated rapidly across multiple health zones over the next months. Initial diagnostic work-up was complicated by armed conflicts in the region and Ebola-virus-centered diagnostic assumptions [64, 65], delaying identification of BDBV as the causative agent until mid-May [4]. In response, WHO and national authorities rapidly escalated outbreak control measures and initiated a clinical research framework to support evaluation of candidate interventions in the absence of licensed BDBV-specific vaccines or therapeutics. In this context, treatment prioritization is now focused on a limited set of repurposed or investigational options, including MBP-134, Maftivimab, remdesivir, and combination antibody-based approaches [66]. Notably, remdesivir served in our study as a reference compound and showed consistent selectivity across the newly established assays. Although remdesivir yielded limited benefit in the PALM trial during the 2018–2020 EBOV outbreak [23], renewed interest has been supported by more recent in vitro minigenome data demonstrating activity against BDBV and preclinical studies against SUDV [66–68]. Within this context, our BDBVΔVP30 data provides additional, authentic *in vitro* evidence supporting this rationale. From a serological perspective, we recently showed that our neutralization assay developed with EBOVΔVP30 outperformed commonly used pseudotyped systems on a predefined panel of serum samples [69], offering a standardized alternative to live-virus studies conducted in biosafety level-4 laboratories. Building on this framework, extending the same neutralization approach to the BDBVΔVP30 system developed in this study would be a timely next step to assess diagnostic samples and vaccine- or antibody-mediated protection in the ongoing outbreak response.

The development of biologically contained systems for EBOV, SUDV, BDBV, TAFV, and RESTV, together with the related MARV, represents a major advance in filovirus research. Our system panel now includes representatives of the currently known human-infecting filovirus species. In addition, the modular design of these systems should facilitate rapid adaptation to newly emerging strains by introducing relevant sequence changes into the antigenome constructs. Beyond antiviral screening, these tools also provide a flexible framework for studying viral replication dynamics, resistance evolution, and other fundamental aspects of filovirus biology, overall boosting preparedness against current and future filovirus outbreaks.

## Acknowledgements

The authors wish to thank Medicines for Malaria Venture, Evotech and CD3 for providing compounds, and Dr. Piet Maes for his contributions to the conceptualization and initial phases of the project.

## Author contributions

This work was conceived by JV and BV. JV, JSt, JSN, CW, KG and BVH performed experiments. JSc, WC, TF and VL performed high-content analysis. JV, JSN and CW performed data analysis. KV supplied reagents and materials. JV, BV and KV wrote the manuscript. All authors read the manuscript and approved its submission.

## Supplementary data

**Figure S1:**
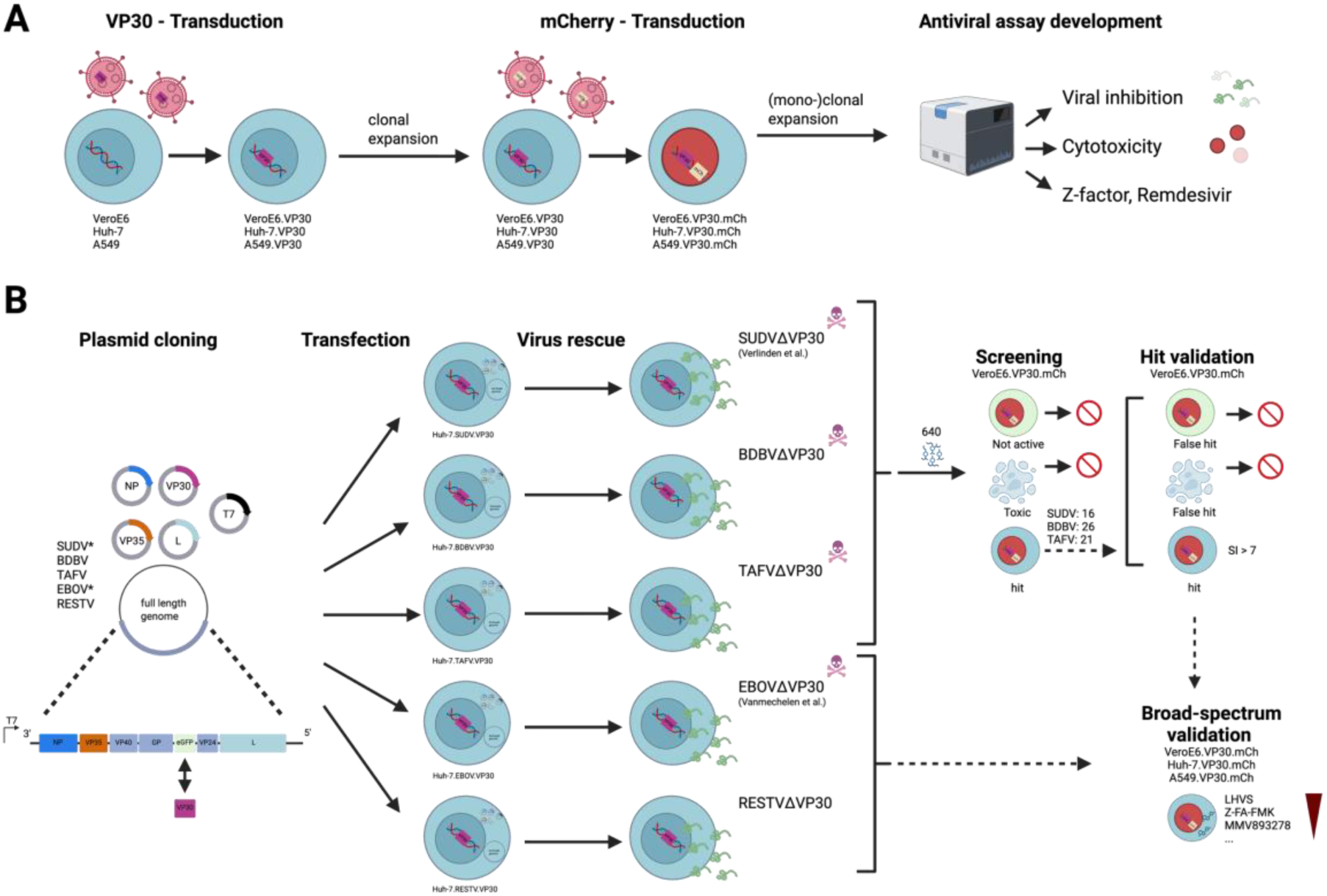
A comprehensive strategy for antiviral discovery against Orthoebolaviruses. The study combines (i) near-completion of a toolkit of biologically contained, known human-pathogenic orthoebolavirus systems, including the addition of RESTV, and (ii) development and application of a dual-reporter antiviral screening platform followed by a compound screening case study. **(A)** Wild-type VeroE6, Huh-7, and A549 cells were sequentially transduced to stably express VP30 (purple), an essential factor required for replication of VP30-deficient viruses, followed by nuclear-localized mCherry (beige, abbreviated as “mCh”) to enable quantification of cell nuclei. Monoclonal selection yielded stable double-transduced reporter cell lines suitable for high-content imaging assays, allowing simultaneous assessment of viral replication (via virus-encoded eGFP; see panel B) and compound-induced cytotoxicity (via mCherry). Assay performance and robustness were validated using remdesivir and Z-factor analysis. **(B)** VP30- deficient antigenomes of BDBV, TAFV, RESTV, SUDV, and EBOV, together with the corresponding support plasmids, were cloned and transfected into VP30-expressing cells to rescue biologically contained viruses, as confirmed by built-in eGFP expression. Primary screening of 640 small molecules (Medicines for Malaria Venture) was performed against SUDV, BDBV, and TAFV in VeroE6 cells, followed by downstream validation across additional Orthoebolaviruses (EBOV and RESTV) and human-derived cell lines to assess broad-spectrum antiviral activity. EBOV and SUDV rescue systems were established previously.

**Figure S2:**
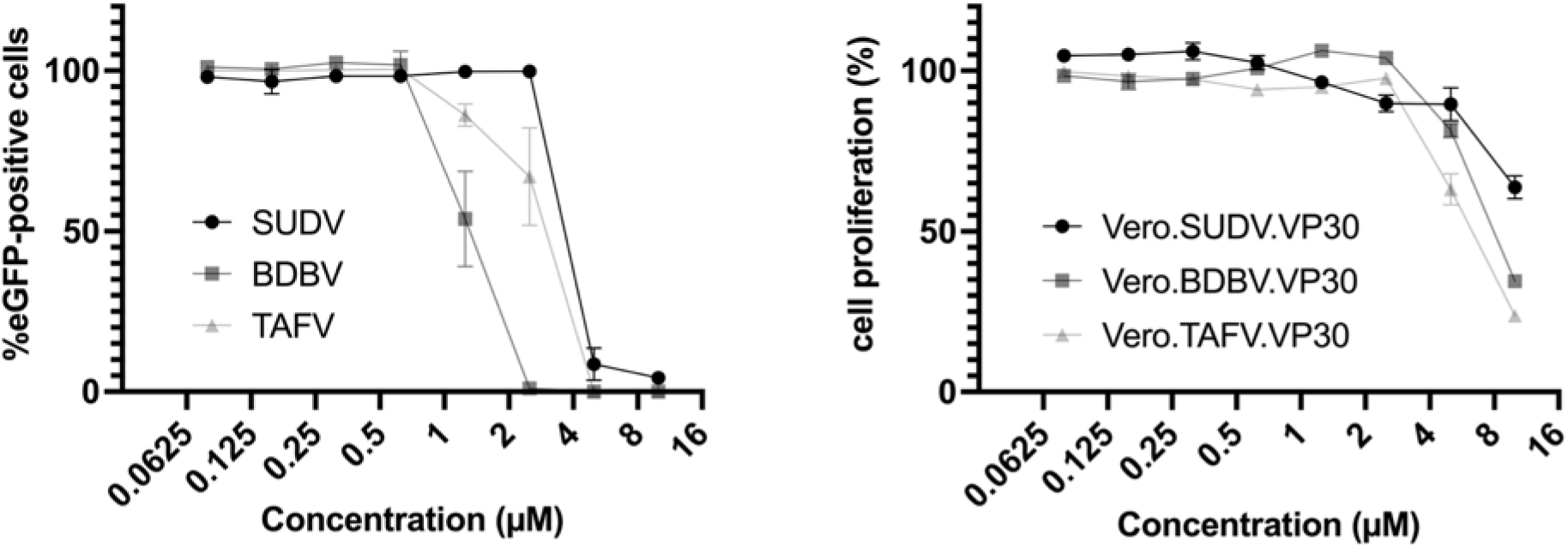
Dose-response curves illustrating the effect of the P-glycoprotein inhibitor CP-100356 against SUDV, BDBV, and TAFV in VeroE6 cells. Antiviral activity was assessed based on the percentage of eGFP-positive infected cells (left). Normalized cell proliferation, serving as a proxy for cytotoxicity, was simultaneously measured by quantifying the total number of nuclei positive for nuclear mCherry fluorescence (right). Data points represent mean values ± standard deviations of six replicates.

**Figure S3:**
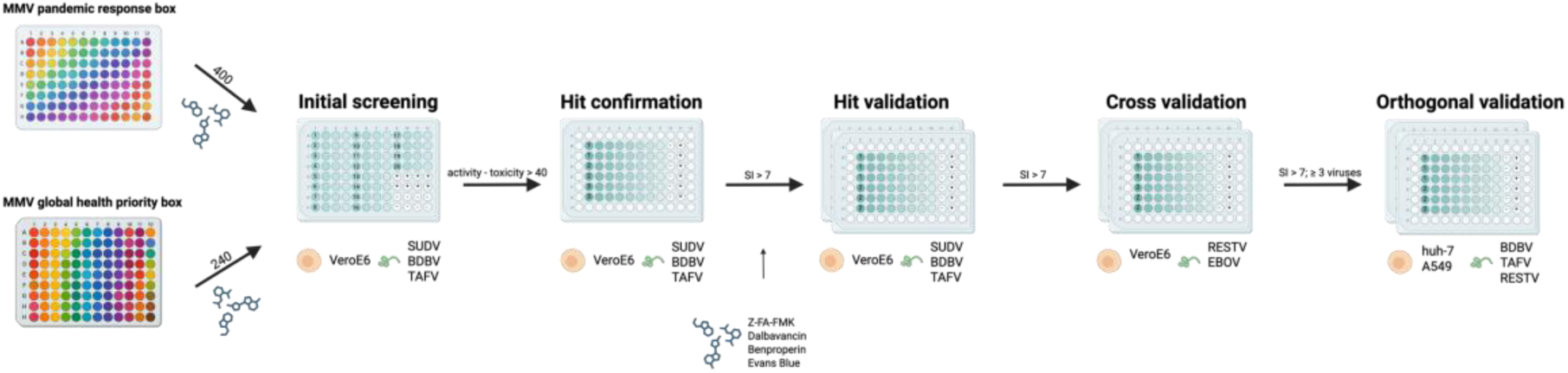
General workflow for compound screening and validation using biologically contained Orthoebolaviruses. A total of 640 compounds from two Medicines for Malaria Venture (MMV) libraries, the Pandemic Response Box (400 compounds) and the Global Health Priority Box (240 compounds), were initially screened against SUDV, BDBV, and TAFV in VeroE6 cells using a four-fold dilution series in single replicate. Compounds displaying antiviral activity exceeding cytotoxicity by more than 40% were subjected to hit confirmation using nine-point dilution series in duplicate. Confirmed hits (SI > 7) were further validated in 3–6 replicates, and 4 compounds active against EBOV from a previous study were included. Cross-validation was performed against RESTV and EBOV for all remaining hits, followed by orthogonal validation in Huh-7 and A549 cells for compounds active against at least three viruses. Virus-only and medium-only control wells were included on each plate for normalization, and only internal wells were used for compound testing, as outer wells were excluded due to edge effects.

**Table S1:** Optimized assay conditions in 96-well format per virus–cell line combination.

| Virus | Cell line | #cells seeded | %FBS | Viral input (log<sub>10</sub> TCID<sub>50</sub>) | Read-out dpi | Z-factor |
| --- | --- | --- | --- | --- | --- | --- |
| BDBV | VeroE6 | 20000 | 2 | 1.1 | 6 | 0.95 |
| BDBV | Huh-7 | 10000 | 10 | * | 6 | 0.50 |
| BDBV | A549 | 7500 | 10 | 0.97 | 10 | 0.93 |
| TAFV | VeroE6 | 20000 | 2 | 1.14 | 5 | 0.98 |
| TAFV | Huh-7 | 7500 | 10 | 0.98 | 4 | 0.96 |
| TAFV | A549 | 10000 | 10 | 1.56 | 6 | 0.92 |
| RESTV | VeroE6 | 20000 | 2 | 1.2 | 7 | 0.95 |
| RESTV | Huh-7 | 12500 | 10 | 1.03 | 5 | 0.92 |
| RESTV | A549 | 10000 | 10 | 0.61 | 8 | 0.96 |
| SUDV | Huh-7 | 10000 | 8 | * | 4 | 0.28 |
| SUDV | A549 | 10000 | 10 | * | 6 | 0.29 |

The table summarizes the assay setup used for each virus across VeroE6, huh-7, and A549 cell lines. Parameters include the number of cells seeded per well, the percentage of fetal bovine serum (FBS) supplemented in the medium, the viral input expressed as log<sub>10</sub> TCID<sub>50</sub>, the selected day post-infection (dpi) for fluorescence-based read-out and associated Z-factors. \* indicates that the viral titer could not be determined, and consequently, the undiluted virus stock was directly applied to the cells.

